# Jumping DNA polymerases in bacteriophages

**DOI:** 10.1101/2024.04.26.591309

**Authors:** Natalya Yutin, Igor Tolstoy, Pascal Mutz, Yuri I Wolf, Mart Krupovic, Eugene V Koonin

## Abstract

Viruses with double-stranded (ds) DNA genomes in the realm *Duplodnaviria* share a conserved structural gene module but show a broad range of variation in their repertoires of DNA replication proteins. Some of the duplodnaviruses encode (nearly) complete replication systems whereas others lack (almost) all genes required for replication, relying on the host replication machinery. DNA polymerases (DNAPs) comprise the centerpiece of the DNA replication apparatus. The replicative DNAPs are classified into 4 unrelated or distantly related families (A-D), with the protein structures and sequences within each family being, generally, highly conserved. More than half of the duplodnaviruses encode a DNAP of family A, B or C. We showed previously that multiple pairs of closely related viruses in the order *Crassvirales* encode DNAPs of different families. Here we identify four additional groups of tailed phages in the class *Caudoviricetes* in which the DNAPs apparently were swapped on multiple occasions, with replacements occurring both between families A and B, or A and C, or between distinct subfamilies within the same family. The DNAP swapping always occurs “in situ”, without changes in the organization of the surrounding genes. In several cases, the DNAP gene is the only region of substantial divergence between closely related phage genomes, whereas in others, the swap apparently involved neighboring genes encoding other proteins involved in phage replication. We hypothesize that DNAP swapping is driven by selection for avoidance of host antiphage mechanisms targeting the phage DNAP that remain to be identified, and/or by selection against replicon incompatibility. In addition, we identified two previously undetected, highly divergent groups of family A DNAPs that are encoded in some phage genomes along with the main DNAP implicated in genome replication.

## Introduction

Viruses with large double-stranded (ds) DNA genomes in the realm *Duplodnaviria* share a uniformly conserved structural gene module but vary greatly in their repertoires of DNA replication proteins (Koonin et al., 2020a; Weigel and Seitz, 2006). Some viruses encode most of the proteins required for DNA replication, whereas others rely (almost) entirely on the replication machinery of the host. Generally, the self-sufficiency of DNA replication correlates with the viral genome size. DNA polymerases (DNAPs) are central components of the viral replication systems that are present in more than half of the available genomes of duplodnaviruses greater than 40kb in size (Kazlauskas et al., 2016). There are four major DNAP families involved in the genome replication in cellular life forms, families A, B, C and D (hereafter PolA-D), with the A, B and C families also being common among DNA viruses. The core catalytic domains of these DNAPs adopt three unrelated folds, namely, (i) the RNA Recognition Motif (RRM), often called the Palm domain (joins the accessory Thumb and Fingers domains) in PolA and PolB, (ii) nucleotidyltransferase Polβ-like fold in PolC, and (iii) the double-psi beta-barrel domain in PolD (Czernecki et al., 2023; Kazlauskas et al., 2020; Koonin et al., 2020b; Raia et al., 2019; Sauguet, 2019). In bacteria, PolC is the primary polymerase responsible for the genome replication, whereas PolA is involved in DNA repair processes; PolB is rare in bacteria and is apparently derived from viruses (Kazlauskas et al., 2020; Kornberg and Baker, 1992). In archaea, replication is catalyzed by either PolB or PolD, and paralogs of PolB are also involved in repair (Greci and Bell, 2020). In eukaryotes, almost all processes of DNA synthesis involved in both replication and repair are catalyzed by DNAPs of the PolB family in the nucleus and PolA in mitochondria (Burgers, 2009; Burgers and Kunkel, 2017).

Different groups of tailed viruses of the class *Caudoviricetes* infecting bacteria and archaea encode PolA, PolB or PolC (or no DNAP at all), PolA being the most common, and PolC the rarest (Kazlauskas et al., 2016). All large dsDNA viruses of eukaryotes, in the realms *Duplodnaviria* (phylum *Peploviricota*) and *Varidnaviria* (phylum *Nucleocytoviricota*), and unassigned class *Naldaviricetes* (baculo-like viruses), employ PolB (Kazlauskas et al., 2020). Many smaller viruses (with <50 kb genomes), especially in the realm *Varidnaviria* (e.g., polintons, adenoviruses, tectiviruses), replicate with the help of a distinct variety of B family DNAPs, the protein-primed PolB (Kazlauskas et al., 2016; Krupovic and Koonin, 2015; Nasko et al., 2018; Schoenfeld et al., 2013), or, less commonly, PolA, also referred to as TV-Pol (Iyer et al., 2008). Notably, archaeal viruses exclusively encode family B DNAPs which can be either RNA- or protein-primed (Makarova et al., 2014; Prangishvili et al., 2017).

The DNAPs are essential proteins that are highly conserved within each family, at the sequence and structure levels (Kazlauskas et al., 2016; Raia et al., 2019). Therefore, it came as a surprise that among closely related genomes of phages in the order *Crassvirales*, multiple replacements of PolA with PolB and vice versa were observed (Yutin et al., 2021). Similar replacement of replication proteins was detected also among smaller phages of the order *Vinavirales* (Krupovic and Bamford, 2007; Yutin et al., 2022). It is particularly notable that in each of these cases, the replacements occurred within otherwise conserved genomic contexts.

In this work, we aimed to systematically identify and explore cases of between-family DNAP swapping in *Caudoviricetes*. We show that DNAPs were swapped repeatedly in the evolution of multiple groups of tailed phages.

## Methods

### Dataset of phage genomes and phage genome tree

Genome-wide relationships between the 18,382 *Caudoviricetes* genomes, available in GenBank as of November 2022, were analyzed using reciprocal best hits between viral protein sequences as follows. The set of non-nested ORFs of at least 75 bp was obtained for each virus genome using the NCBI ORFfinder tool (https://www.ncbi.nlm.nih.gov/orffinder/). Reciprocal best hits for all pairs of genomes (*A*,*B*), covering at least 50% of the query sequences were identified between the two ORF complements using BLASTP (Altschul et al., 1997). Distance between the two genomes was calculated as

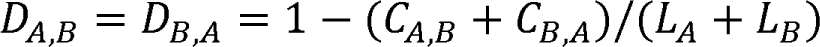

where *C*_A,B_ is the total length of the part of genome *A*, covered by ORFs that have reciprocal best hits in genome *B* and *L*_A,_ is the length of genome *A* (ditto for *C*_B,A_ and *L*_B_).

The tree was reconstructed from the pairwise distance matrix using the FastMe 2.0 program (Lefort et al., 2015) and ultrameterized by iteratively balancing subtrees, descending from each internal node.

### Identification of phage DNA polymerases

PolA, PolB, PolC reference protein sequences were collected from the NCBI virus database (https://www.ncbi.nlm.nih.gov/labs/virus/vssi/#/) and from the respective publications, in particular, the PolA sequences were from (Dorawa et al., 2022; Keown et al., 2022; Nasko et al., 2018); PolB sequences were from (Kazlauskas et al., 2020), and PolC sequences were from (Timinskas and Venclovas, 2019) (https://ftp.ncbi.nih.gov/pub/yutinn/jumping_polymerases_2024/). Open reading frames (ORFs) from the 18,382 *Caudoviricetes* genomes were searched for polymerases using BLASTP with collected reference PolA, PolB, PolC proteins as queries (e-value threshold of 0.0001). The initial set of hits was clustered using MMSEQS2 (Steinegger and Soding, 2017) at similarity threshold 0.5; sequences within clusters were aligned using MUSCLE5 (Edgar, 2022). Cluster alignments were iteratively compared to each other using HHSEARCH and aligned using HHALIGN (Zimmermann et al., 2018). Sequences in the final alignments were examined for the presence of the critical catalytic residues. Full-length sequences predicted to be catalytically active were retained for the downstream analysis (see Supplementary Table S1 for the final set of polymerases, classified as active).

DNAP swapping hotspots were identified by the presence of DNAPs from different families within a subtree of depth 0.15 (corresponding to ∼1/3rd of the total tree depth). Sister subtrees exhibiting polymerase diversity were grouped into DNAP-swapping clades.

### Comparison of phage genomes

Pairwise genome alignments were constructed using Mauve (Darling et al., 2004) and visualized using with Geneious Prime® 2022.1.1 (https://www.geneious.com). Predicted phage proteins were annotated using CDD (Marchler-Bauer et al., 2017) and HHPRED (Soding, 2005; Zimmermann et al., 2018).

### Phylogenetic analysis of phage proteins

The identified viral PolA, PolB, PolC sequences were combined with homologs identified in a collection of completely sequenced bacterial and archaeal genomes downloaded from NCBI Genomes (https://ftp.ncbi.nlm.nih.gov/genomes/ASSEMBLY_REPORTS/) in November 2021. Sequences of phage DNAPs of the order *Crassvirales* were added from (Yutin et al., 2021). The protein sequences were aligned using MUSCLE5 (Edgar, 2022). Phylogenetic trees were constructed using IQ-TREE 2 (Minh et al., 2020), with the following models chosen according to BIC by the built-in model finder: VT+F+R10 for PolA, Q.pfam+F+R8 for PolB, and VT+F+R5 for PolC, and visualized with MEGA11 (Tamura et al., 2021).

Large terminase subunits were aligned using MUSCLE5 (Edgar, 2022); constrained and unconstrained phylogenetic trees were reconstructed using IQ-TREE 2 (Minh et al., 2020) with the automatically selected evolutionary models and compared using the built-in Approximately Unbiased test.

### Protein structure prediction and analysis

MSAs for divergent family A DNA polymerases identified in this study (divPolA1 and divPolA2) were submitted to a local installation of ColabFold (colabfold_batch with default settings except “–num-models 1 –num-recycle 3”) (Mirdita et al., 2022). In addition, all individual divPolA1 and divPolA2 were modeled with a singularity version of AlphaFold2 (Jumper et al., 2021) (version 2.2.0 with the following specifications: “--db_preset=full_dbs –model_preset=monomer_ptm –max_template_date=2022-10-01”) on the high performance cluster BIOWULF at the NIH. All models were compared to a local version of pdb70 (created on December 10, 2021) using Dali (Holm, 2020) to identify closest structures. Structure-guided alignments between representative divPolAs and closest related structures were obtained using the Dali web server (Holm, 2022), and key residues were identified. Representative structures modeled with AlphaFold2 (divPolA1 clade5: CAB4155247, clade6: CAB4155247 and divPolA2 AUR84708) were displayed and superimposed with the respective DNAP structures from pdb using ChimeraX (Pettersen et al., 2021).

### Data Availability

This work is based on the analysis of genomes publicly available in GenBank. All other data generated by this analysis is contained in the Supplementary Material or publicly available at https://ftp.ncbi.nih.gov/pub/yutinn/jumping_polymerases_2024/

## Results

### DNA polymerase diversity in *Caudoviricetes*

In the analyzed set of 18,382 *Caudoviricetes* genomes, we identified 6560 PolA, 2857 PolB, and 947 PolC proteins (Supplementary Table S1). The *Caudoviricetes* genome tree was split into subtrees at the depth of 0.15, roughly corresponding to a genus level (see an example in Supplementary Figure S1; https://ftp.ncbi.nih.gov/pub/yutinn/jumping_polymerases_2024/genome_tree/). Of the 1,514 subtrees that included more than one virus genome, 563 were found to encode a DNAP, and 8 encoded more than one DNAP. Analysis of the DNAP distribution pattern revealed two distinct types of DNAP heterogeneities within phage subtrees: (i) DNAPs of different families were encoded in closely related viruses, with a single copy in each genome, implying swapping of DNAP genes (6 subtrees), and (ii) the same viral genome encoded two DNAPs of different families (2 subtrees).

### DNA polymerase swapping in *Caudoviricetes*

We identified 6 subtrees in the phage genome tree in which the phages encoded DNAPs of different families. Representative pairs of most closely related genomes with different family DNAPs from these subtrees are listed in Table 1. To investigate the provenance of these groups of phages encoding distinct DNAPs, we examined deeper clades in the phage genome tree that included each of the 6 subtrees (Table 1), apart from *Crassvirales* for which frequent PolA-PolB swaps have been reported previously (Yutin et al., 2021). Clades 1, 2, and 3 consisted of PolA- and PolB-encoding genomes, and clade 4 genomes encompassed PolA and PolC (Figures 1, 2). The DNAPs from these clades were reciprocally mapped onto the corresponding PolA, PolB, and PolC trees (Supplementary Figure S2). Examination of this mapping suggests that, in addition to inter-family DNAP swaps, some intra-family DNAP swaps occurred in the selected clades, that is, PolA or PolB was apparently replaced by a distinct DNAP of the same family on several occasions. Below we discuss each of these clades in detail, in an attempt to reconstruct the evolutionary scenarios.

**Figure 1.**
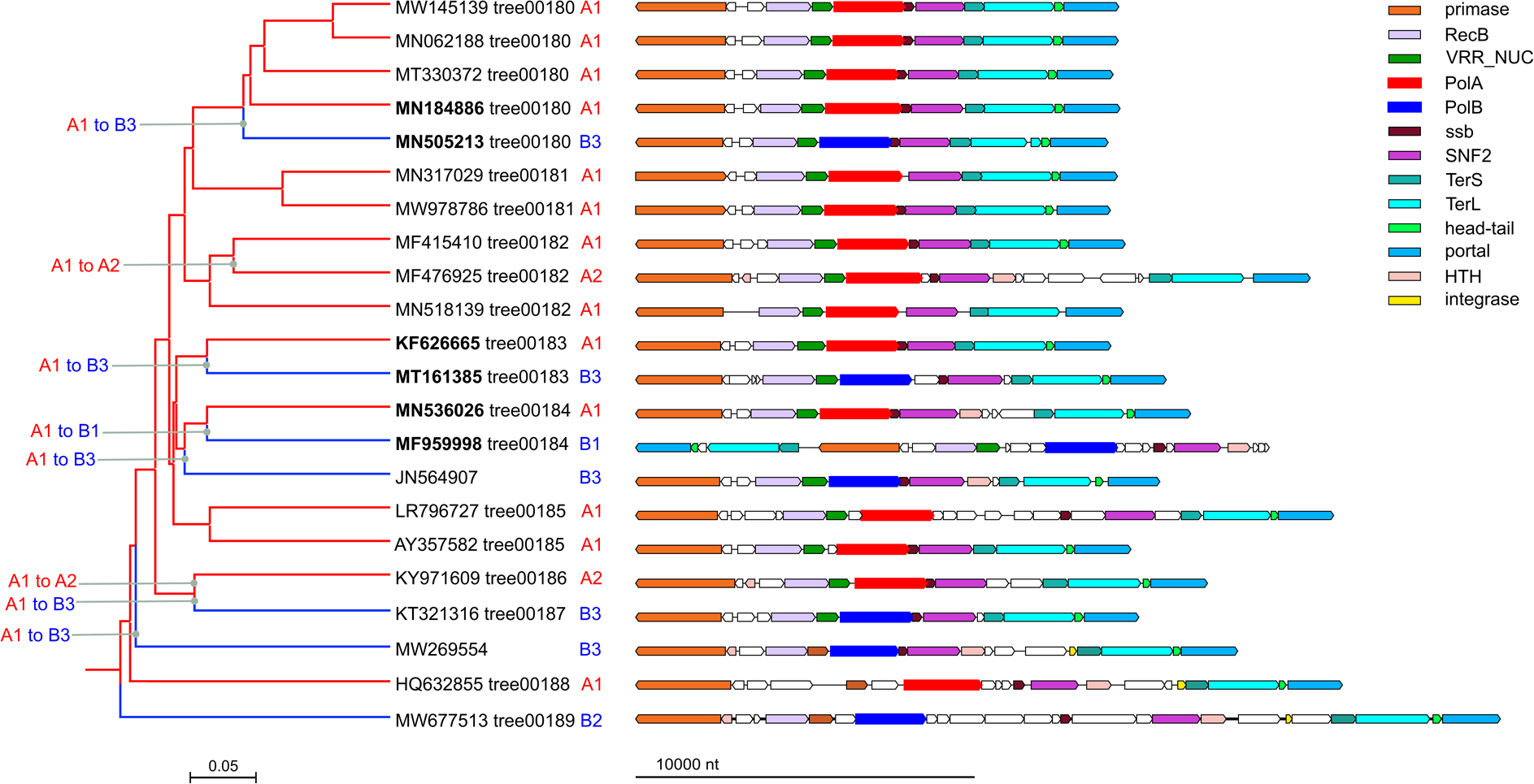
DNA polymerase swapping in Clade 1. *Left*: Clade 1 genome tree, reduced to salient representatives. DNAP families and clades are marked on tree leaves; tree edge colors indicate the polymerase families; inferred DNAP swapping events are marked on the corresponding tree edges. *Right*: genome maps of polymerase neighborhoods; homologous genes are shown in the same colors.

**Table 1.** Phage clades displaying DNAP swapping and examples of exchange between closely related genomes DNAP genome Organism genome

| *DNAP* | *genome* | *Organism* | *genome length* | *ANI or AAI* | *clade* |
| --- | --- | --- | --- | --- | --- |
| tree00180 |  |  |  |  |  |
| PolA | MN184886.1 | Erwinia phage pEp_SNUABM_08 | 62716 | 39% AAI | Clade1; subtrees 00180-00189 |
| PolB | MN505213.1 | Serratia phage JS26 | 63971 |  |  |
| tree00183 |  |  |  |  |  |
| PolA | KF626665.1 | Phage Sano | 56147 | 39% AAI | --“-- |
| PolB | MT161385.1 | Xanthomonas phage FoX4 | 60418 |  |  |
| tree00184 |  |  |  |  |  |
| PolA | MN536026.1 | Pseudomonas phage vB_Pae-SS2019XI | 57567 | 38% AAI | --“-- |
| PolB | MF959998.1 | Marinobacter phage PS6 | 58226 |  |  |
| tree01348 |  |  |  |  |  |
| PolA | MZ477002.1 | Acinetobacter phage Phab24 | 93604 | 92.72% ANI | Clade2; subtrees 01336-01351 |
| PolB | MN276049.1 | Acinetobacter phage BS46 | 94068 |  |  |
| tree01419 |  |  |  |  |  |
| PolA | MG592602.1 | Vibrio phage 1.237.B._10N.261.52.C5 | 60160 | 99.54% ANI | Clade3; subtrees 01419-01425 |
| PolB | MG592464.1 | Vibrio phage 1.089.O._10N.261.51.F9 | 59851 |  |  |
| tree01472 |  |  |  |  |  |
| PolA | MT601273.1 | Bacillus phage vB_BsuS-Goe12 | 124287 | 99.74% ANI | Clade4; subtrees 01466-01481 |
| PolC | MT601274.1 | Bacillus phage vB_BsuS-Goe13 | 126848 |  |  |

We sought to validate the intra-clade cross-family swaps of DNAPs using the large subunit of the terminase (TerL) as the reference. The TerL sequences were collected from the genomes with identified DNAPs within each of the clades 1, 2, 3 and 4 and clade-specific TerL phylogenetic trees were constructed. Then, we constructed topologically constrained trees, separating the TerL from genomes encoding different DNAPs (for example, for clade 1, the constraint separated TerL from PolA- and PolB-bearing genomes). Such constrained topologies represent hypothetical phylogenies where DNAPs of different families are not intermixed within the clade histories. The optimal TerL trees satisfying these constraints were compared to the unconstrained trees, in an attempt to falsify the scenario with multiple DNAP swaps. For all clades, the Approximately Unbiased test decisively rejected the constrained topologies (Supplementary Table S2), suggesting that multiple DNAP swaps within each clade did occur.

Clade 1 included 126 genomes from several genera of the family *Casjensviridae* (genome size range 50-70 kb) of which 96 encoded PolA whereas the remaining 30 encoded PolB. The genome tree for this clade is dominated by a distinct group of PolA (A1 in Figure 1) in which 3 disjointed PolB branches (B1-B3) and two branches from a separate group of PolA (A2) are embedded. Altogether, comparison of the phage genome tree with the phylogenetic trees of PolA and PolB suggests 8 independent DNAP swaps including both 6 inter-family (PolA to PolB) exchanges and 2 intra-family (A1 to A2) exchanges (Figure 1). In most of the phage genomes in this clade, the swapped PolA and PolB genes share the same or similar genomic neighborhood (Figure 1).

Clade 2 (subtrees 01336-01351 in the phage genome tree) unites phages from genera *Plaisancevirus*, *Saclayvirus*, *Barbavirus*, subfamily *Ounavirinae*, and several unclassified *Caudoviricetes* (genome size range 80-105 kb). The phages in subtree 01347 encode an additional protein with remote sequence and structural similarity to PolA, which we discuss in the next section. Here we address the apparent PolA to PolB swaps in this clade (Figure 2). PolBs of Clade 2 are monophyletic (B4 in Figure 2), but PolAs come from three distinct branches (A3, A4 and A5 in Figure 2; see Supplementary Figure S2 for the DNAP trees). As in Clade 1, comparison of the phage genome tree with PolA and PolB phylogenetic trees suggests several independent swaps although the direction and the order of these events is difficult to establish.

**Figure 2.**
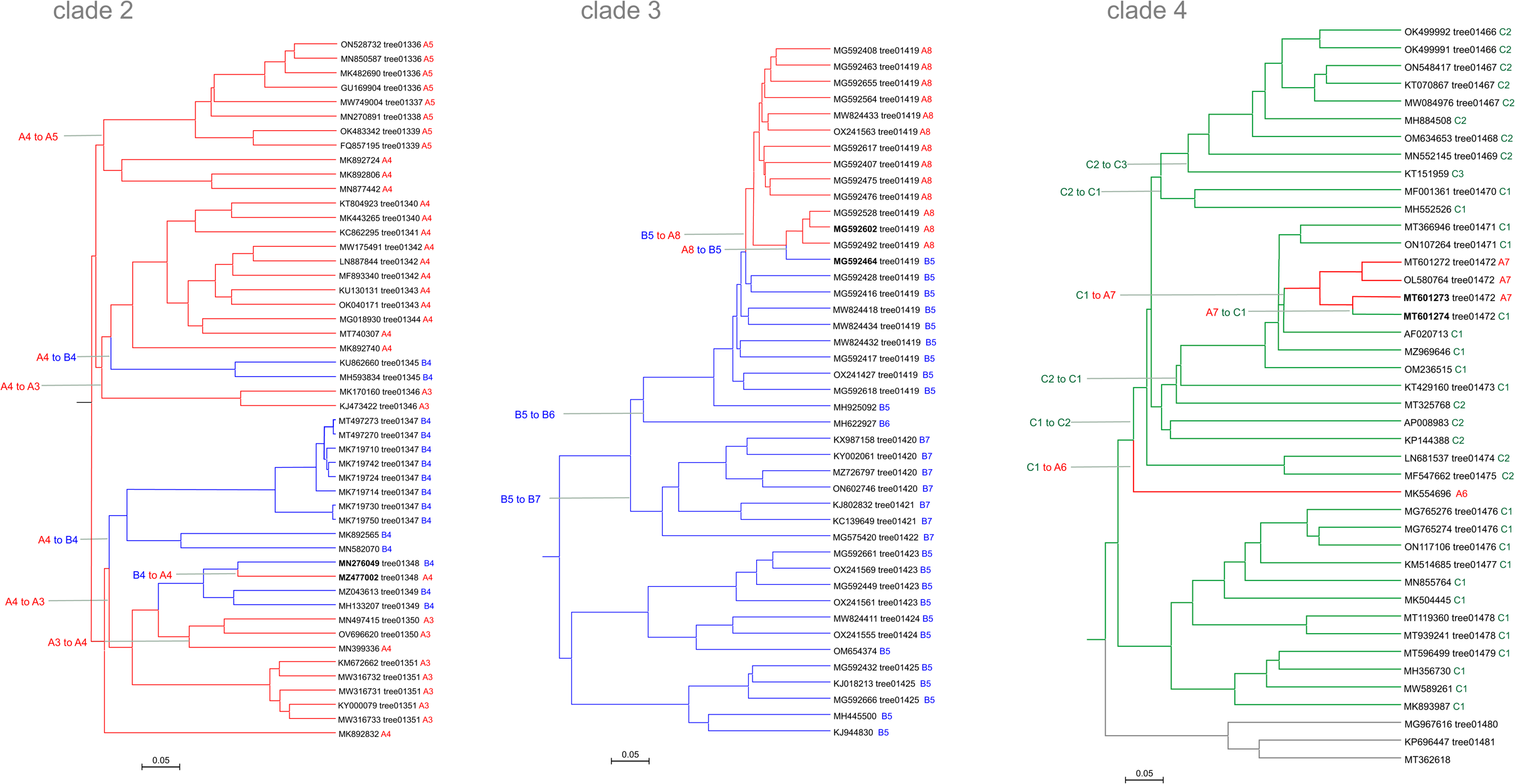
DNA polymerase swapping in Clades 2, 3 and 4. Genome trees of Clades 2, 3 and 4, reduced to salient representatives. DNAP families and clades are marked on tree leaves; tree edge colors indicate the polymerase families; inferred DNAP swapping events are marked on the corresponding tree edges.

Two Acinetobacter phages of Clade 2, Phab24 (MZ477002) and BS46 (MN276049), have average nucleotide identity (ANI) of 93%, and yet, encode DNAPs of different families, PolA and PolB, respectively (Table 1; Figure 3a). Comparison of the genome organization in the vicinity of the DNAP genes (in which we additionally included the corresponding genome region of Acinetobacter phage TaPaz (MZ043613) because its PolB is most similar to PolB of MN276049 (Acinetobacter phage BS46). suggests that, in this case, the primase-helicase gene was replaced along with the DNAP.

**Figure 3.**
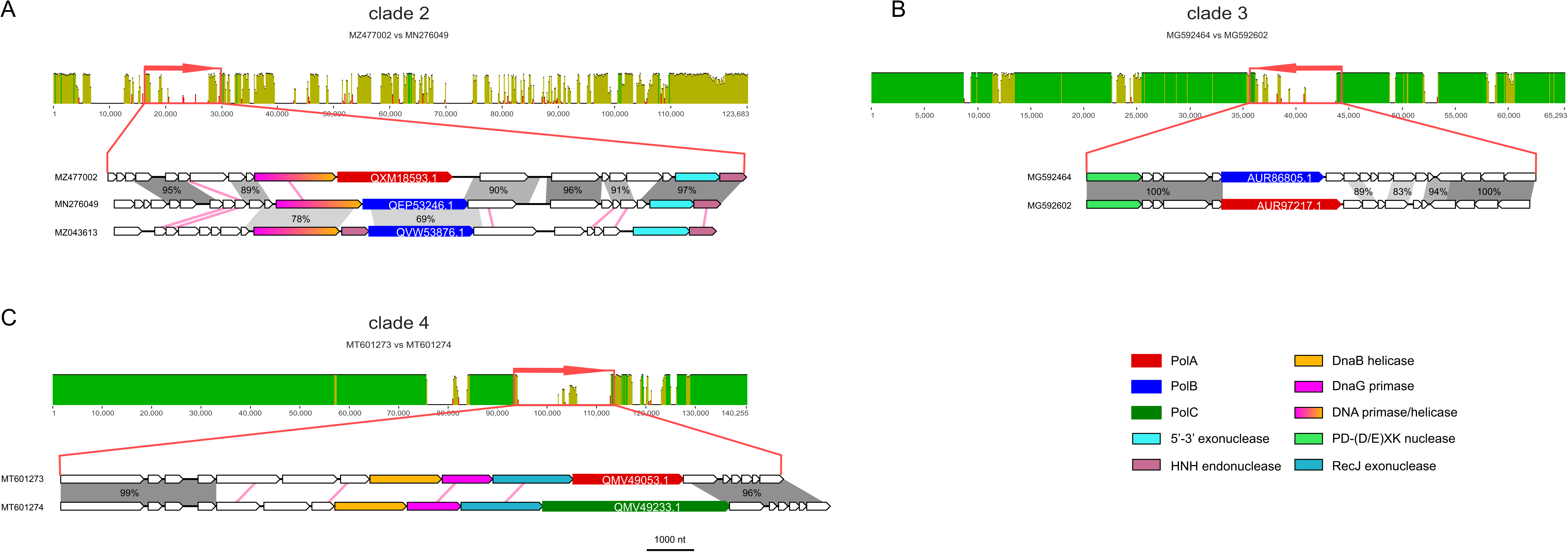
Pairwise alignments of closely related viral genomes encoding DNAPs of different families. A. Clade 2, tree01348 (unclassified Caudoviricetes). MN276049 is permuted at 27000; MZ477002 is reversed. B. Clade 3, tree01419 (unclassified Caudoviricetes). C. Clade 4, tree01472 (Spbetavirus). In the upper part of each panel, nucleotide sequence similarity between the compared genomes is shown in green or yellow, on the scale from 0 to 100%. Red arrows denote the genomic regions containing the DNAP genes, visualized on the genome maps. The lower parts, show the blow up of the regions containing the DNAP genes. Functionally annotated homologous genes are shown in the same colors. DNAP genes are labeled with their GenBank protein IDs. Grey shading highlights genomic regions with high nucleotide sequence similarity (percent identity indicated). Pink lines connect genes with significant detectable amino acid sequence similarity detected with BLASTP but no significant nucleotide sequence similarity.

Clade 3 unites subtrees 01419-01425 (Figure 2) and includes phages from genera *Sashavirus*, *Nonanavirus*, *Gorganvirus*, and unclassified *Caudoviricetes*, with genome size range of 42.5-62.5 kb. In this clade, a single PolB to PolA swap appears to have occurred within the subtree01419, whereas the basal PolB genes belong to distinct clades (Figure 2). In this clade, we identified a pair of nearly identical genomes, Vibrio phage 1.237.B._10N.261.52.C5 (MG592602) and Vibrio phage 1.089.O._10N.261.51.F9 (MG592464), that encode different family DNAPs, PolA and PolB, respectively. Pairwise genome comparison shows that the DNAP neighborhood is the only large region with markedly lower similarity between the two genomes (Figure 3b). The DNAP gene is the only one that obviously was replaced but additional rearrangements might have occurred in the adjacent genomic region containing genes encoding uncharacterized small proteins (Figure 3b).

Clade 4 unites subtrees 01466-1481 including families *Tybeckvirinae* and *Andrewesvirinae*, genera *Audreyjarvisvirus*, *Spbetavirus*, *Latrobevirus*, *Sextaecvirus*, *Slashvirus*, and several unclassified *Caudoviricetes* (genome size range 61-185 kb). In this clade, the phages encode either PolC or PolA. PolC, specifically group C1, is likely to be ancestral in this assemblage of phages (Figure 2 and Supplementary Figure S2). This ancestral PolC apparently was replaced with PolA on two independent occasions (Figure 2), and furthermore, underwent several intra-family replacements involving PolC variants from groups C2 and C3. We also identified a pair of closely related genomes in this clade with over 99% ANI, Bacillus phage vB_BsuS-Goe12 (MT601273) and Bacillus phage vB_BsuS-Goe13 (MT601274), that encode DNAPs of different families, PolA and PolC, respectively. Comparative genome analysis of this pair of phages revealed an extended segment of dissimilarity suggesting that several replicative genes, including DnaB-like helicase, DnaG-like primase and RecJ-like exonuclease, traveled together with the DNAPs, but the replacement keeps the gene context unchanged, that is, the genes appear to have been replaced *en bloc* (Figure 3c).

When the inter-family swaps were found to have occurred on shallow branches of the virus tree, these events typically involved phages that shared bacterial hosts (Supplementary Figure S3).

### Two novel phage PolAs

In addition to the typical DNAPs, we identified two divergent variants of PolA encoded in phage genomes. One of these, denoted divPolA1, is present in a group of 14 phages (Flavobacterium phage vB_FspM_immuto_3-5A and related phages), with genomes in the range of 155-190 kb that also encode a ‘regular’ DNAP, either family A, B, or C (Figure 4a). The conserved arrangement of genes implicated in replication downstream of the divPolA1 gene suggests that divPolA1 is involved in replication (Figure 4b). Structural prediction for divPolA1 (Figure 5) showed that it contains a Palm domain in which the main catalytic residues are conserved, whereas the Thumb domain is truncated and the 3’ exonuclease domain seems to be inactivated, with all three catalytic aspartates replaced (Figure 5). Thus, divPolA1 most likely retains the DNAP activity whereas the exonuclease activity is lost.

**Figure 4.**
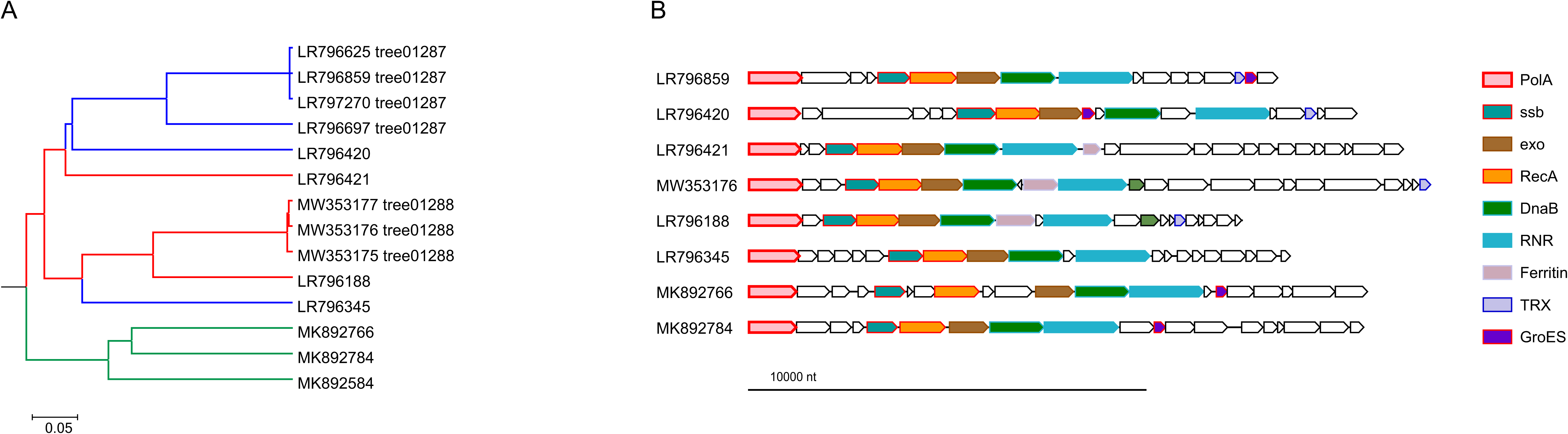
Phylogenetic and genomic context of divPolA1. A, Genome tree of divPolA1-encoding viruses. Tree branches are marked according to the identity of the ‘regular’ polymerases: red, PolA; blue, PolB; green, PolC. B, divPolA1 genome neighborhoods; homologous genes are shown in the same colors. Genomes: LR796625, LR796859, LR797270, LR796697, LR796420, LR796421, LR796188, LR796345: uncultured Caudovirales phages; MW353177: Flavobacterium phage vB_FspM_immuto_13-6C; MW353176: Flavobacterium phage vB_FspM_immuto_3-5A; MW353175: Flavobacterium phage vB_FspM_immuto_2-6A; MK892766: Prokaryotic dsDNA virus sp. isolate GOV_bin_1807; MK892784: Prokaryotic dsDNA virus sp. isolate GOV_bin_703; MK892584: Prokaryotic dsDNA virus sp. isolate GOV_bin_630.

**Figure 5.**
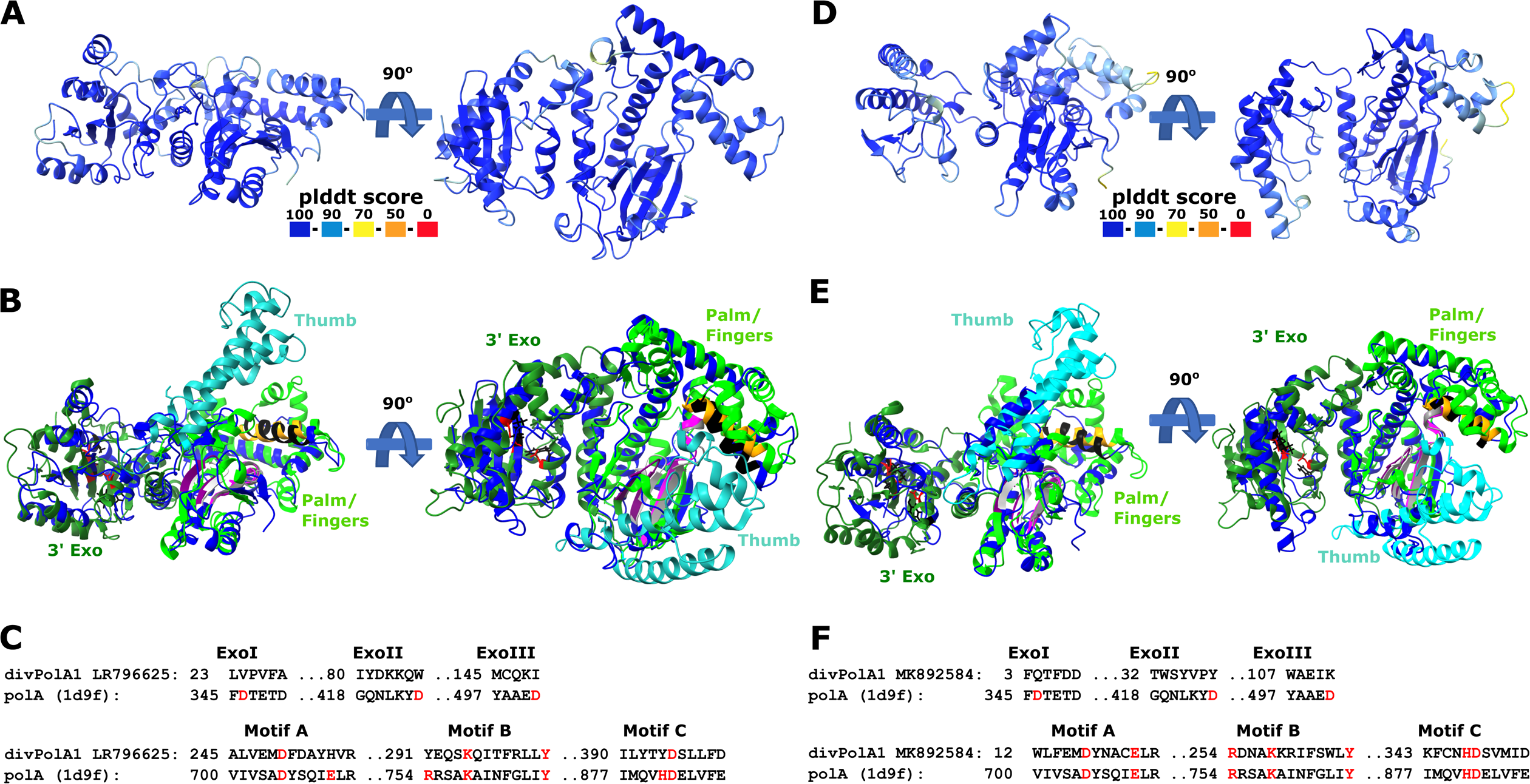
divPolA1 structure prediction. A, D. Predicted representative divPolA1 structures colored according to plddt score (AlphaFold2 model; A: CAB4155247, genome ID: LR796625; D: QDP51333, genome ID: MK892584). B, E. Structural comparison of divPolA1 (blue) and a representative DNA polymerase I (pdb 1d9f, Klenow fragment). The representative DNAP I is colored by domain organization: 3’ exonuclease domain (3’ Exo, dark green), thumb domain (cyan) and palm/finger domain (green). Sites of motifs A, B and C highlighted in magenta/grey, orange/black and purple/light grey for DNAP I and divPolA1, respectively. Aspartic acid residues in 3’ exonuclease motifs I, II and III of DNAP I are highlighted in red, the corresponding sites in divPolA1 in black. C, F. Structure-guided alignments of selected 3’ exonuclease and palm/finger domain motifs between divPolA1 representative (C: CAB4155247, genome ID: LR796625; F: QDP51333, genome ID: MK892584) and PolA 1D9F_A. Key residues highlighted in red.

Another diverged PolA variant, divPolA2, was initially identified in 110 genomes of barbaviruses (Rheinheimera phage vB_RspM_Barba18A and related viruses) with genomes of 80-85 kb, each also encoding a regular PolB. Additional PSI-BLAST searches against the *Caudoviricetes* database using barbavirus divPolA2s as queries revealed more divPolA2 proteins, sometimes with two or three paralogs per phage genome (Figure 6). Unlike the PolB of these phages, which is embedded within a typical context of replication-related genes, the gene encoding divPolA2 is located in variable gene neighborhood. Structural modeling suggests that divPolA2 contains an active DNAP (Palm) catalytic domain but lacks a Thumb domain homologous to those of any other DNAPs (Figure 7). Instead, this protein contains an N-terminal globular domain without detectable similarity to any other known domains that potentially might function as the Thumb. As in the case of divPolA1, the 3’ exonuclease domain is lacking. Most likely, divPolA2 is not the replicative enzyme of barbaviruses, a role that belongs to PolB. Instead, divPolA2 might be a DNAP involved in repair processes, or an RNA polymerase, given that PolA was co-opted for that function in T7 and related phages (Czernecki et al., 2023). Of note, structural comparison did not only reveal DNAPs as the top hits for divPolA2, but also DNA-directed RNA polymerases (mitochondrial RNA polymerase (PDB ids: 7a8p, 6ymv, Dali z-score ∼12) and, with lower z-score (∼9), also a viral DNA-directed RNA polymerase from bacteriophage N4 (genus *Enquatrovirus*, class *Caudoviricetes*) (PDB id: 4ff3). These observations are compatible with the possibility that divPolA2 is actually an RNA polymerase although the N-terminal globular domain of divPolA2 is unrelated to the N-terminal domains of PolA-related RNA polymerases (Figure 7).

**Figure 6.**
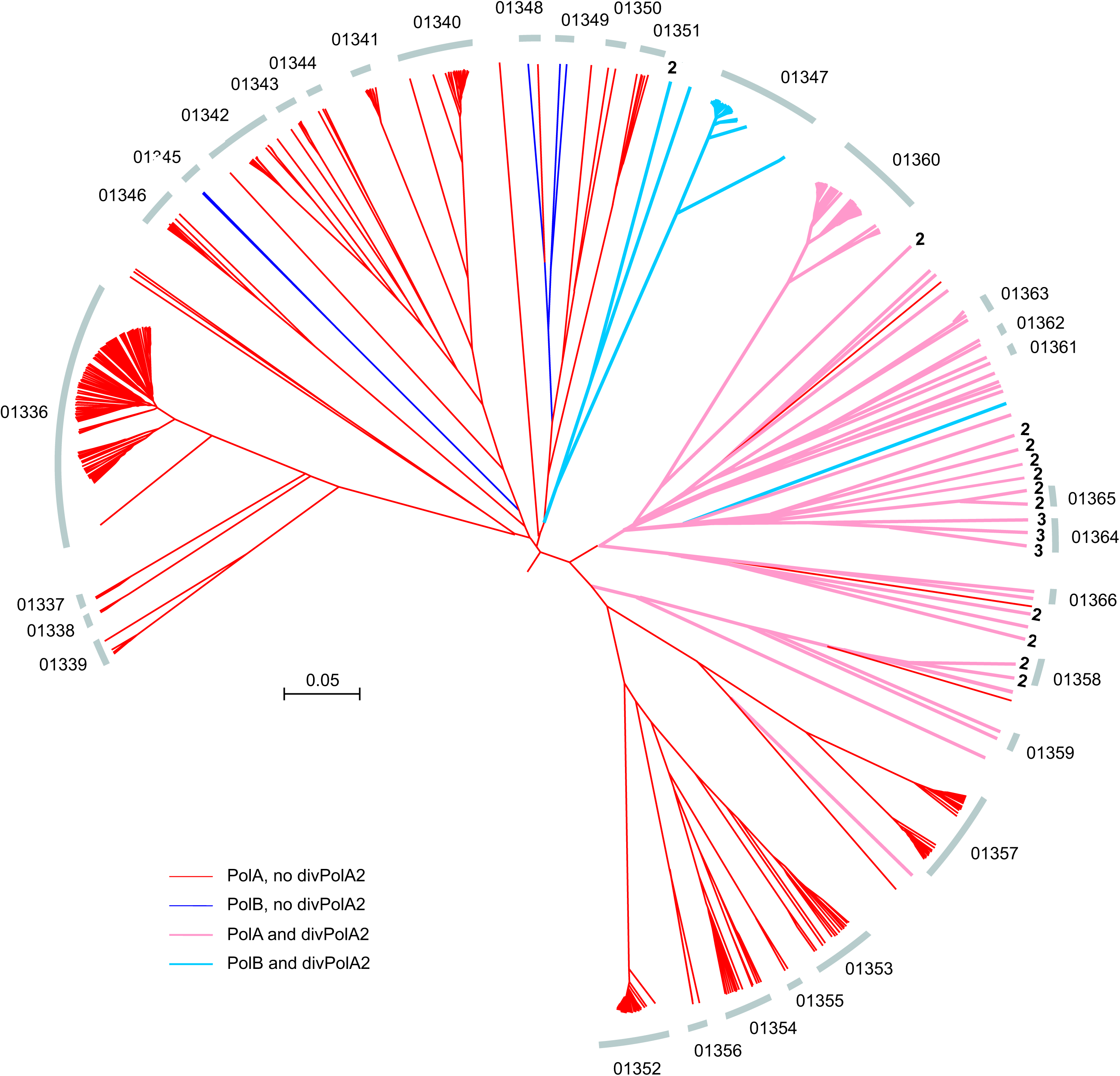
Phylogenetic context of divPolA2. Genome tree for the clade containing divPolA2 genes is shown. Arcs indicate subtrees. Numbers at tree tips indicate the number of divPolA2 paralogs. Barbaviruses are located in the subtree 01347.

**Figure 7.**
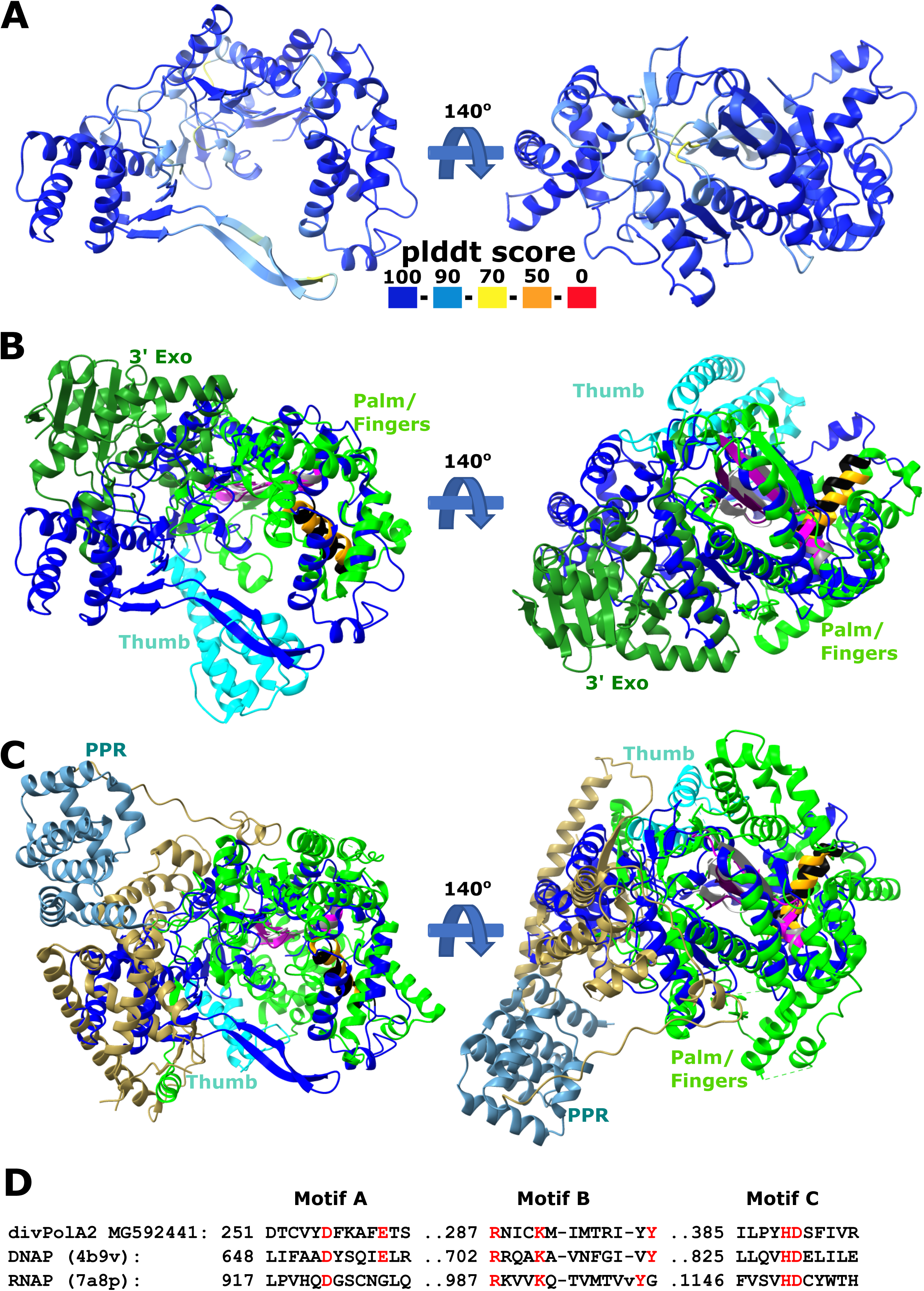
divPolA2 structure prediction. A. Predicted representative divPolA2 structure colored according to plddt score (AlphaFold2 model; AUR84708, genome MG592441.1). B,C. Structural comparison of divPolA2 (blue) and a representative DNA polymerase I (pdb 4b9v, B) and a representative RNA polymerase (pdb 7a8p, human mitochondrial RNAP, C). The representative DNAP I and RNAP are colored by domain organization: 3’ exonuclease domain (3’ Exo, dark green, DNAP I only), thumb domain (cyan) and palm/finger domain (green) and RNAP N-terminal pentatricopeptide domain (PPR, cornflower blue). Sites of motifs A, B and C highlighted in magenta/grey, orange/black and purple/light grey for divPolA2 and DNAP I, respectively. D. Structure-guided alignments of selected motifs of palm/finger domain between divPolA2 and DNAP I 4b9v and RNAP (7a8p). Key residues highlighted in red.

## Discussion

Tailed viruses of bacteria and archaea that comprise the class *Caudoviricetes* in the realm *Duplodnaviria* are considered to be the most abundant group of viruses on earth (Mushegian, 2020; Suttle, 2005). Although the virion structures and the core structural proteins are conserved throughout the realm, these viruses greatly differ in their genome size and gene repertoires. In particular, some caudoviricetes encode a (nearly) complete suite of proteins required for replication, whereas others have none, and the entire range of intermediates exists as well (Kazlauskas et al., 2016). This variety notwithstanding, more than half of the caudoviricetes encode a DNAP – the obvious centerpiece of the replication machinery – that belongs to either A or B, or C family. Generally, the DNAP is a conserved component of the replication apparatus. Unexpectedly, however, in our previous comparative genomic analysis of *Crassvirales* (the order of *Caudoviricetes* that includes the most abundant viruses identified in the human gut), we found that DNAPs were swapped between closely related phages on multiple occasions, with PolB replacing PolA or vice versa (Yutin et al., 2021). Intrigued by this observation, we probed a much broader range of phages and report here that multiple DNAP swaps occurred in at least four additional phage groups.

The DNAP replacements involved either different families, that is, PolA to PolB and vice versa, as well as PolC to PolA, or distinct groups within the same DNAP family. Remarkably, these replacements in each case occurred “in situ”, without a change in the neighboring gene arrangement. The swap involved either the DNAP gene alone or several adjacent genes encoding other components of the replication machinery, but in each case, the gene replacement appears to have occurred with “surgical precision”. The genes for proteins involved in replication tend to cluster in viral genomes (Kazlauskas et al., 2016), and the preservation of their order upon DNAP swapping implies that coregulation of these genes is important for phage reproduction.

The recurrent DNAP swapping in phage evolution raises intriguing questions on both the molecular mechanisms of these exchanges and the selective forces that could drive them. The mechanisms of DNAP swapping remain enigmatic considering the striking precision of these events. Whether or not the phages involved in the swaps are within the range of sequence identity required for homologous recombination, it hardly can contribute to the capture of distantly related genes. Whether the replacing DNAP comes from a prophage integrated in the host cell genome or a coinfecting phage, illegitimate recombination seems to be essential, and the positive selection associated with the swap should be strong enough to provide for the fixation of the rarely emerging precise replacements. In cases where we could pinpoint the intra-family DNAP swaps to a narrow phylogenetic context (that is, between closely related phages), they typically occurred between phages that infect the same host (at least up to the genus level; Supplementary Figure S3). These observations are compatible with the involvement of coinfection, either cotemporaneous or sequential, in DNAP exchanges.

With respect to the evolutionary forces driving DNAP swapping, it has been shown that multiple defense systems specifically target the phage replication machinery components (Stokar-Avihail et al., 2023). Involvement of at least four types of known defense mechanisms can be suspected. Mutations within DNAP genes have been demonstrated to allow phages to escape restriction by the poorly understood Borvo defense system (Millman et al., 2022; Stokar-Avihail et al., 2023). Although the mechanism of Borvo activation remains unclear, it has been suggested that the DNAP structure, its complex with other proteins and/or DNA encompasses molecular patterns that activate Borvo (Huiting and Bondy-Denomy, 2023). Similarly, AbiQ, a type III toxin-antitoxin abortive infection system, was shown to be activated by various phage proteins, with escape mutants localized to a family A DNAP (Samson et al., 2013). Another recent study has similarly shown that mutations in PolB of T-even phages enabled escape from DarTG, a type II toxin-antitoxin system that provides immunity by ADP-ribosylating phage DNA (LeRoux et al., 2022). Furthermore, pattern recognition systems, in particular, those centered at antivirus STAND ATPases (Avs), have been shown to target conserved viral structural proteins, such as the terminase large subunit and the portal protein (Gao et al., 2022; Kibby et al., 2023). These viral proteins are conserved at the level of structures even if their sequences diverge relatively fast. The DNAPs, although not universal among tailed phages, unlike terminase and portal, are common and even more highly conserved at the sequence level, and therefore, the existence of multiple pattern recognition systems targeting DNAPs appears likely.

The high sequence conservation of the DNAPs within each family suggests potential involvement of another type of defense, namely, adaptive immunity mediated by CRISPR systems, and more specifically, primed adaptation (Jackson et al., 2019; Shiriaeva et al., 2022). CRISPR spacers targeting conserved sequences in the DNAP genes are likely to retain complementarity level sufficient for primed adaptation longer than they do in the case of less conserved genes, facilitating acquisition of immunity to the respective phages. Furthermore, existence of yet unknown defense mechanisms targeting DNAPs remains a possibility. An additional or alternative driver of replication module swapping between phages could be the incompatibility of closely related replicons within a coinfected cell, analogous to plasmid incompatibility (Igler et al., 2022; Pilosof, 2023). Further study of the notable but not yet well understood phenomenon of DNAP swapping in phages has the potential to reveal unknown facets of interactions between phages and their bacterial hosts as well as conflicts among different phages.

## Conclusions

We show in this work that replacement of DNAPs by distantly related or even unrelated ones is common in the evolution of tailed phages of the class *Caudovirecetes*. Remarkably, DNAP swapping always occurs “in situ”, with the organization of the surrounding genes, typically, encoding other proteins involved in phage genome replication being preserved, whether the DNAP gene is the only region of substantial divergence between closely related phage genomes, or the replacement involves several neighboring genes. We hypothesize that although illegitimate recombination is required for replacement of the DNAP genes, selection driving such replacements is strong enough to allow the rare emerging variants with precise insertion of the new sequence to be fixed in the phage population. The factors underlying this selection likely include avoidance of host defense mechanism, such as Borvo, pattern recognition or CRISPR primed adaptation, that target DNAPs. In addition to DNAP swapping, we identified two previously undetected, highly divergent groups of family A DNAPs that are encoded in some phage genomes along with the main DNAP implicated in genome replication.

## Acknowledgements

This work utilized the computational resources of the NIH HPC Biowulf cluster (http://hpc.nih.gov). N.Y., P.M., Y.I.W. and E.V.K. are supported by the Intramural Research Program of the National Institutes of Health (National Library of Medicine).

