## Supplementary figures and images for "Jumping DNA polymerases in bacteriophages"

### Supplementary figure 1

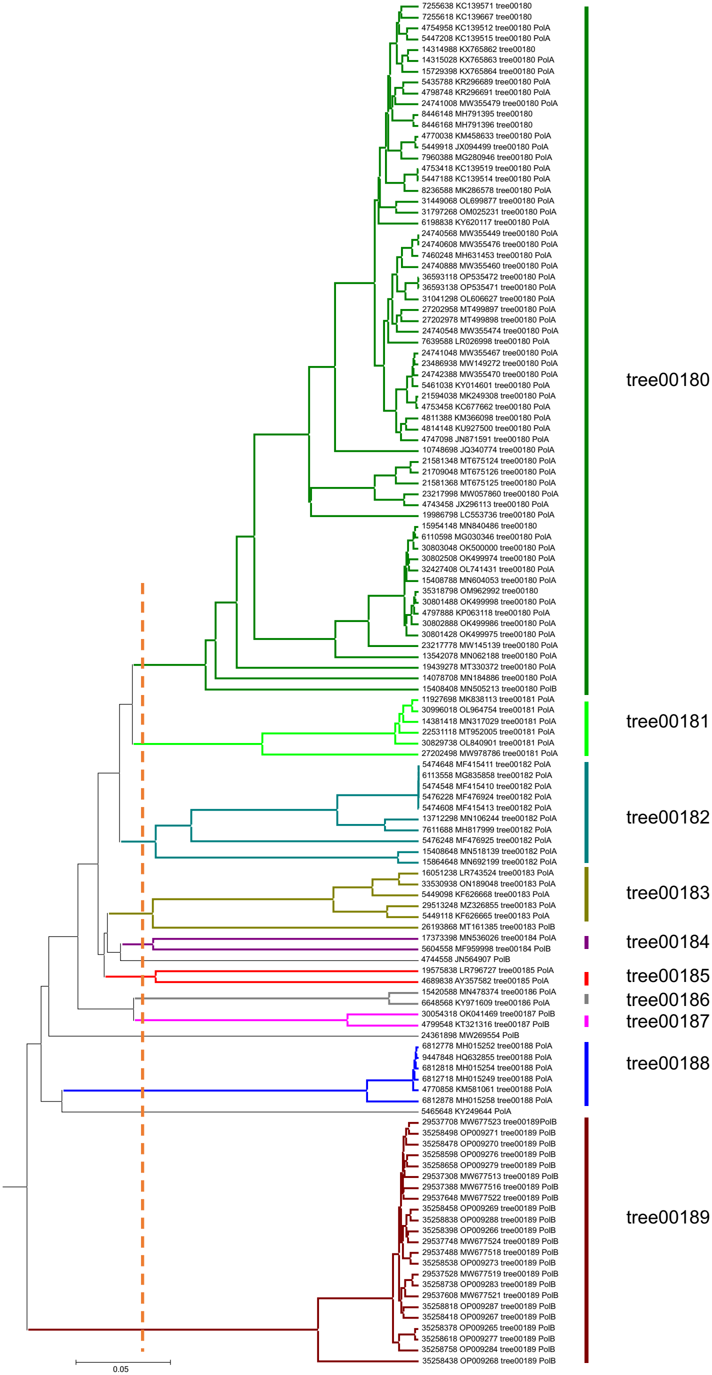

### Supplementary figure 2

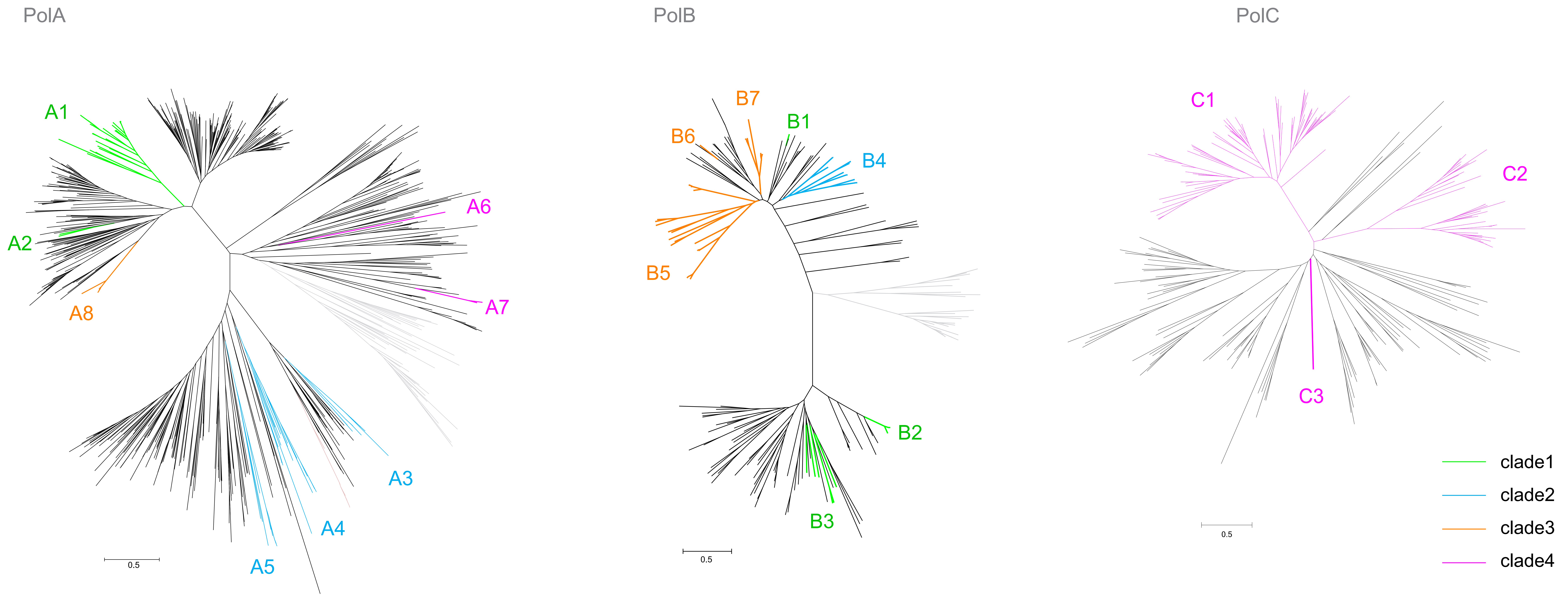

### Supplementary figure 3

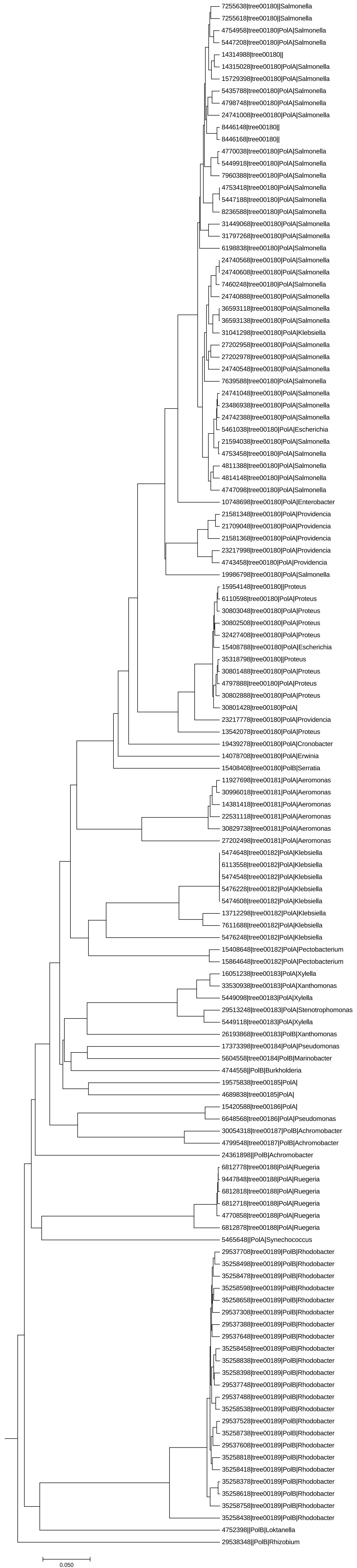

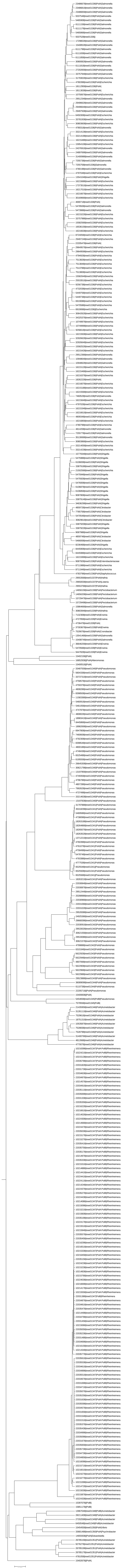

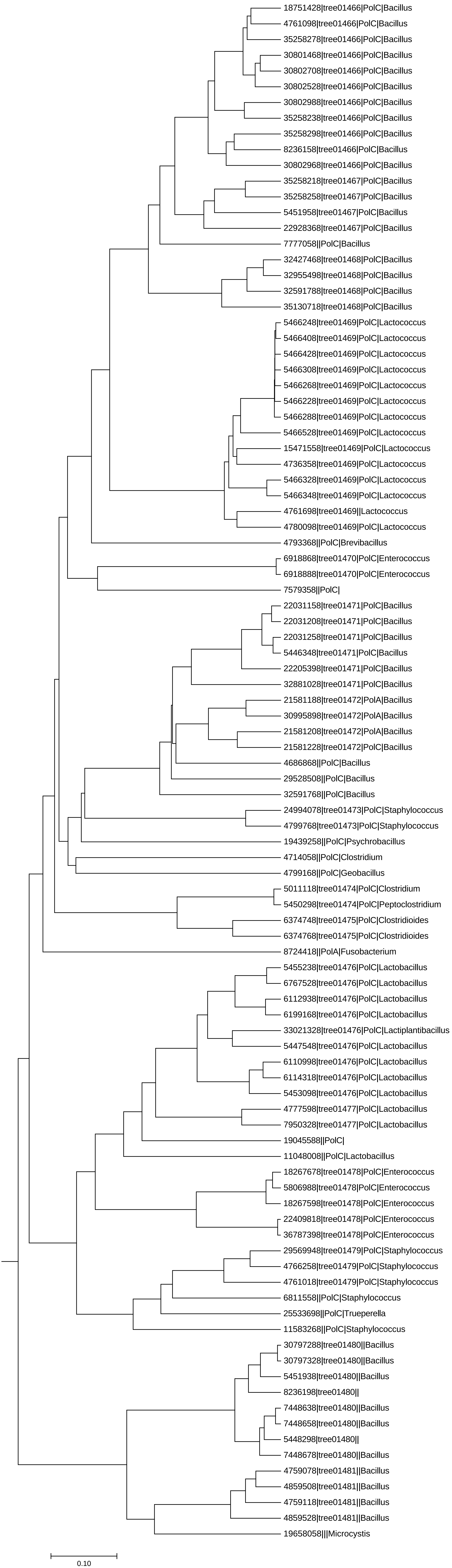

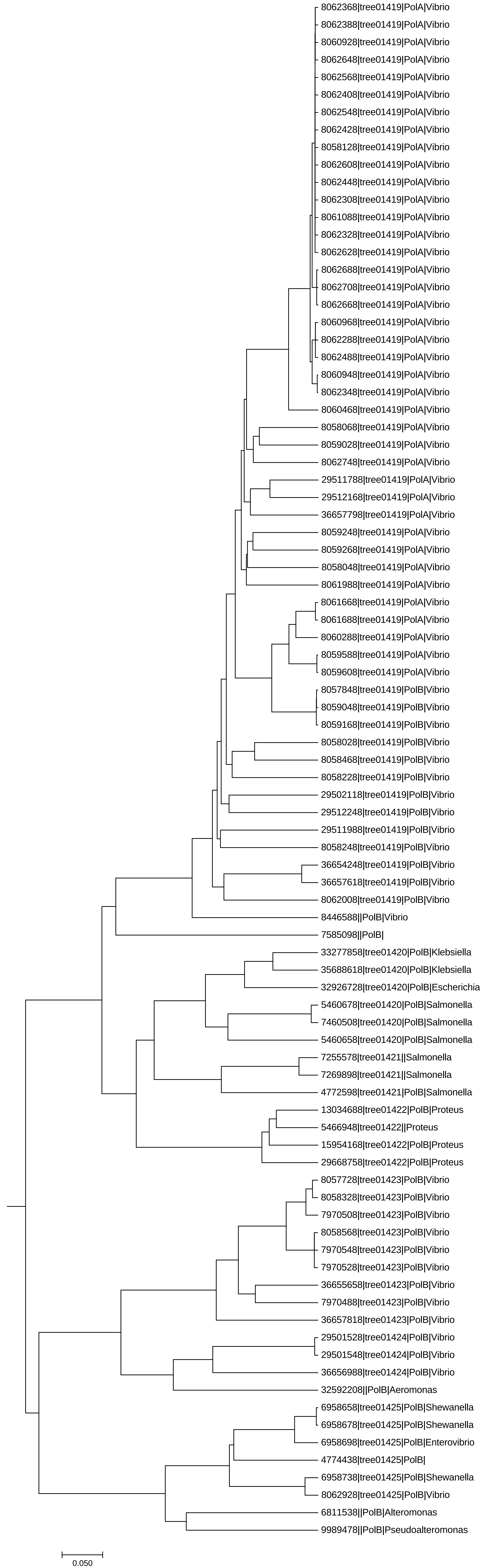
